# Cold Atmospheric Helium Plasma in the Post-Covid Era: A Promising Tool for the Disinfection of Silicone Endotracheal Prostheses

**DOI:** 10.1101/2023.12.10.570744

**Authors:** Diego Morais da Silva, Fellype do Nascimento, Noala Vicensoto Milhan, Maria Alcionéia Carvalho de Oliveira, Paulo Francisco Guerreiro Cardoso, Daniel Legendre, Fabio Gava Aoki, Konstantin Georgiev Kostov, Cristiane Yumi Koga-Ito

## Abstract

The COVID-19 pandemic resulted in a high prevalence of laryngotracheal stenosis. The endoluminal tracheal prostheses used to treat this condition are made of medical-grade silicone (MGS). Despite their excellent properties, the main limitation of these prostheses is the formation of a polymicrobial biofilm on their surfaces that interacts with the underlying mucosa, causing local inflammation and interfering with the local healing process, ultimately leading to further complications in the clinical scenario. Cold atmospheric plasma (CAP) shows antibiofilm properties on several microbial species. The present study evaluated the inhibitory effect of CAP on multispecies biofilms grown on MGS surfaces. In addition to the MGS characterization before and after CAP exposure, the cytotoxicity of CAP on immortalized human bronchial epithelium cell line (BEAS-2B) was evaluated. The aging time test reported that CAP could temporarily change the MGS surface wetting characteristics from hydrophilic (80.5°) to highly hydrophilic (< 5°). ATR-FTIR shows no significant alterations in the surficial chemical composition of MGS before and after CAP exposure for 5 min. A significant log reduction of viable cells in mono-species biofilms (log CFU/mL) of *C. albicans, S. aureus*, and *P. aeruginosa* (0.636, 0.738, and 1.445, respectively) was detected after CAP exposure. Multi-species biofilms exposed to CAP showed significant viability reduction for *C. albicans* and *S. aureus* (1.385 and 0.831, respectively). The protocol was not cytotoxic to BEAS-2B. It could be concluded that CAP can be a simple and effective method to delay the multi-species biofilm formation inside the endotracheal prosthesis.

## 1. INTRODUCTION

Tracheal stenosis is a clinical condition characterized by a reduction of the central airway diameter by a congenital or acquired pathological process. The COVID-19 pandemic has produced a high number of patients submitted to intubation and prolonged mechanical ventilation worldwide, resulting in higher percentages of laryngotracheal stenosis ^1–4^. Commonly, patients with tracheal stenosis after COVID cannot undergo tracheal resection since the maturation of the stenosis, as well as the reduction of inflammation needs to be achieved before the procedure ^5^. Additionally, the recurrence of tracheal stenosis is a risk during the first three months after the surgery ^6^. The prevalence of post-intubation tracheal stenosis in retrospective studies ranges from 6% to 20% and post-tracheostomy from 0.6% to 21% ^7^.

Patients with benign tracheal stenosis or tracheal tumors who are not eligible, either temporarily or permanently, for definitive treatment by tracheal surgical resection are considered for airway stenting to maintain airway patency. Silicone airway stents are widely used for their safety, patient tolerance, easy handling, and lower cost than self-expanding stents. The silicone T-tube was introduced in 1965 ^8^. It requires a tracheostomy for anchoring the horizontal limb of the prosthesis that is occluded by a removable silicone cap. The straight-studded silicone stent was introduced in 1990 ^9^ and does not require a tracheostomy. Both are used as endoluminal support for the trachea, enabling breathing and phonation through the natural airway, thus improving the quality of life ^10^.

Medical grade silicone (MGS) is the primary material in endotracheal prostheses. MGS has attractive properties such as chemical inertness, water and temperature resistance, biocompatibility, and flexibility ^11^. The constant contact between the inhaled air passing through the airway prosthesis and the tracheal underlying mucosa promotes the accumulation of biofilms composed of bacteria and fungi. The interaction between the polymicrobial biofilm and the surrounding mucosa can potentially aggravate the stenosis severity ^12–14^. *Staphylococcus aureus, Pseudomonas aeruginosa*, and *Candida albicans* are the main microorganisms in the polymicrobial biofilm formed inside the patient’s silicone prosthesis implant ^15^. Biofilm formation usually takes approximately seven days, and severe complications such as pneumonia or sepsis can occur ^16^. On the other hand, the MGS degradation promoted by the polymicrobial biofilm is progressive, requiring the prosthesis to be changed every 6 to 12 months after implantation. This can negatively impact the patient’s quality of life and induce higher costs ^10,17^.

Previous studies focused on modifying the MGS surface to promote antimicrobial activity. Silver ions and nanoparticles were used as an alternative approach and were effective against *Escherichia coli, P. aeruginosa*, and *Staphylococcus aureus*. The limitations of these methods are the potential cytotoxicity and mucus accumulation on the silver-coated surface, which can reduce its antimicrobial properties ^18^.

Cold atmospheric plasma (CAP) stands for a low temperature (< 40□C) gas discharge plasma with extensive applications in medical and biomedical fields ^19–22^. The literature has already reported microbial inactivation due to the reactive oxygen and nitrogen reactive species (RONS) and UV radiation produced by CAP. The antimicrobial effect was observed on several species of bacteria and fungi, including *S. aureus, P. aeruginosa*, and *C. albicans* ^23–25^. CAP can be delivered through long, flexible plastic tubes ^26,27^, enabling clinical use. Moreover, previous studies reported that CAP has low toxicity to mammalian cells ^28,29^.

This pre-clinical *in vitro* study focused on the effect of the Helium-CAP over mono-species and multi-species biofilms grown on the MGS surface in a laboratory setting, mimics what is clinically observed inside the T-tube. CAP treatment’s effects on biofilms’ viability were assessed, along with the possible surface changes on the MGS caused by CAP. Lastly, the cytotoxicity of the protocol was tested using a BEAS-2B immortalized human bronchial epithelium cell line. The motivation of the study is to develop a protocol for CAP treatment that can be applied directly to the external limb of the T-tube during outpatient visits, aiming to control the biofilm proliferation in the lumen of the silicone prosthesis. Such features can prolong the prosthesis’s durability, reduce the interaction between the biofilm and the adjacent mucosa, and enhance the local healing process.

## 2. MATERIALS AND METHODS

The MGS samples were prepared in disks measuring 8 mm in diameter by 2 mm in height. The specimens had the same chemical composition as T-tube implants. They were used to perform the surface characterization of silicone before and after the plasma treatment and the microbiological assays. Before treatment, all samples were washed in an ultrasonic bath by immersion for 10 min in water and 10 min in isopropyl alcohol. After that, another step of water washing was performed for 10 min. Finally, the samples were sterilized in an autoclave for 20 min packed in a medical-grade sheet. The MGS samples were stored in a dry place until the experiments.

### 2.1 Characterization of CAP interactions with the MGS surface

Figure 1 illustrates the configuration of the CAP system used to treat MGS samples. The device was described in ^24^. It consists mainly of a dielectric barrier discharge (DBD) type reactor, composed of a metallic pin electrode placed inside a closed-end quartz tube, which in turn is placed inside a dielectric enclosure. The working gas (helium, 99.2% purity) is fed into the chamber and flushed to the ambient air through a 1-meter long and flexible plastic tube connected to the reactor output (inner and outer diameter equal to 2.0 mm and 4.0 mm, respectively). A 0.5-mm-diam copper wire was installed inside the plastic tube and placed a few millimeters inside the reactor to avoid contact with the quartz tube. The other tip of the copper wire terminates 2 mm before the output tip of the plastic tube. The high voltage is turned on when the working gas flows, and a primary discharge is ignited inside the DBD reactor. The latter polarizes the copper wire, and a small plasma jet is ignited at the end of the plastic tube.

**Figure 1.**
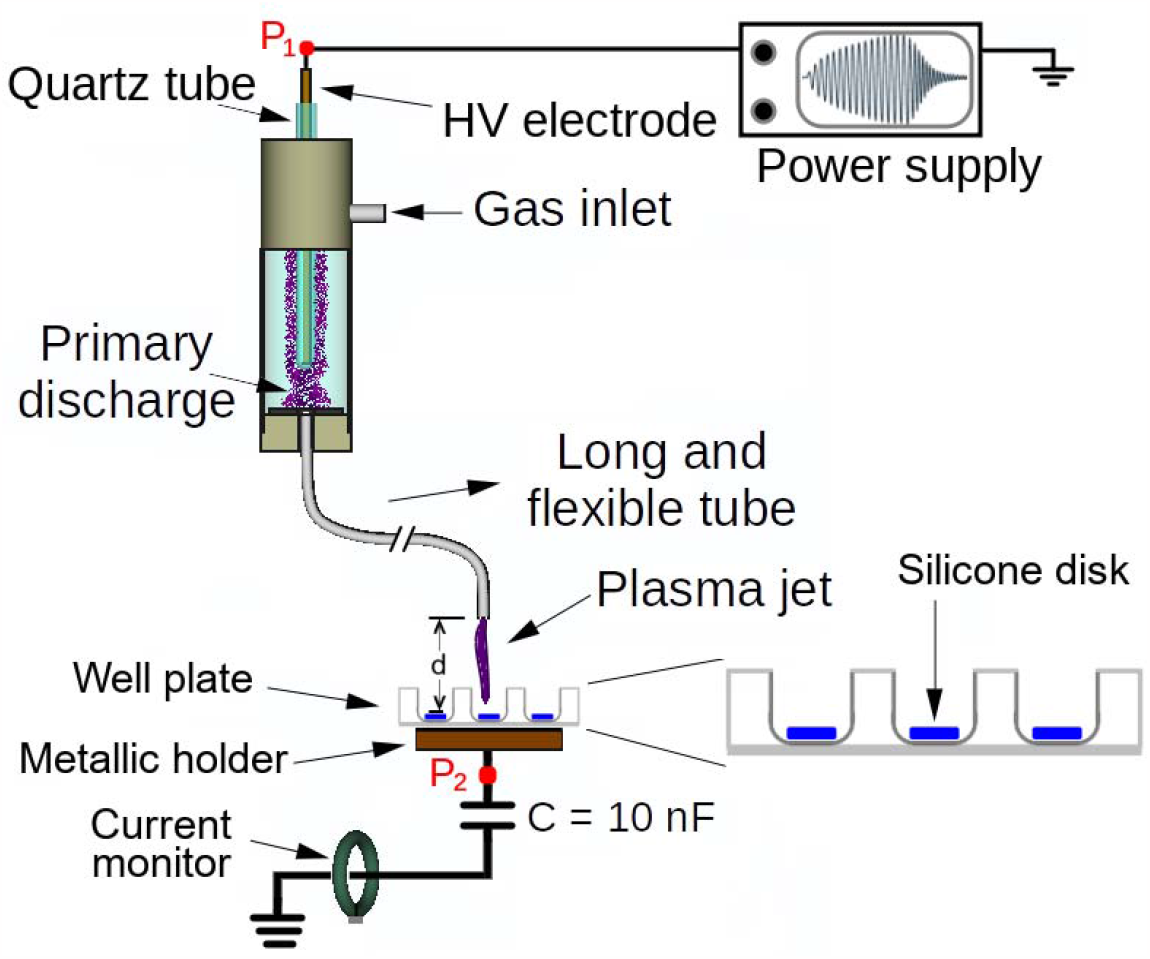
Scheme for CAP generation and MSG treatment.

A commercial AC generator from GBS Elektronik GmbH (model Minipuls4) was used as the power source to generate the CAP. It was used to produce an amplitude-modulated high-voltage (HV) waveform that consists of a sinusoidal “burst” with an oscillation frequency of 31.7 kHz, followed by a voltage-off interval, which repeats at a repetition period (T) equal to 1.2 ms.

MGS samples were placed inside a 24-well plate at a distance d = 5.0 mm from the plasma outlet. The employed gas flow rate was 2.0 SLM. Aiming to study the interaction between the CAP and the MGS, as well as the behavior of the MGS surface after 2 minutes of plasma exposure, the CAP was generated by applying HV with an amplitude of 17.2 kV-peak-to-peak to the pin-electrode, which is nearly 40% higher than the value employed for the treatment of biofilms. This value was chosen to simulate an extreme case and evaluate the possibility of material damage due to plasma exposure. The samples were characterized by wettability measurements and Fourier transform infrared spectroscopy (FT-IR) before and after CAP treatment. To assess the water contact angle (WCA), the samples were analyzed using an F300 Rame-Hart goniometer. Six WCA measurements were performed, one before plasma exposure and five after the treatment (at time instants of 0 min, 15 min, 30 min, one hour, and two hours). Due to the small sample size, different samples were used for each WCA measurement. To analyze possible changes in the surface chemistry of the MGS samples exposed to CAP treatment, FT-IR spectra were recorded before and right after the CAP exposure. Such measurements were carried out using the ATR mode of a Lambda-100 Perkin-Elmer spectrometer in the region between 4000 and 400 cm^-1^, with a 4 cm^-1^ resolution averaged over 16 scans.

### 2.2 Formation of mono-species biofilms on MGS surfaces

Reference strains (Table 1) were plated in Brain Heart Infusion (BHI) agar for bacteria or Sabouraud agar for fungi. Plates were incubated for 24 h at 37 °C under aerobiosis. After, standardized suspensions (1x10^7^ CFU/mL) were prepared in sterile saline (NaCl 0.9%) with the aid of a spectrophotometer (AJX-1600 – AJ Micronal). The optical density and the wavelength adopted for each microorganism are shown in Table 1. Sterile MGS specimens were transferred to a 24-well plate, and 2.0 mL of brain heart infusion (BHI) broth and 200 µL of the microbial inoculum were added to each well. Plates were incubated for 48 h at 37 °C under aerobiosis and agitation (120 rpm). The culture medium was refreshed after 24 h.

**Table 1.** Data on the reference strains used for biofilm formation and parameters of optical density (O.D) and wavelength (λ) adopted for standardized suspension preparation (1x10^7^ CFU/mL).

| Microorganism | Reference number | Optical density | Wavelength (nm) |
| --- | --- | --- | --- |
| *Candida albicans* | ATCC 18804 | 0.760 | 530 |
| *Pseudomonas aeruginosa* | ATCC 27853 | 0.130 | 600 |
| *Staphylococcus aureus* | ATCC 6538 | 0.249 | 490 |

### 2.3 Formation of multi-species biofilms on MGS surfaces

The microbial suspensions were obtained as described in 2.2. The multispecies biofilms were grown on the surface of MGS inside a 24-well plate. 2.0 mL of BHI broth + 2.5% Bovine Serum Albumin (BSA) were added to each well. Then, 200 µL of the *C. albicans* and *S. aureus* inoculum were added, followed by 20 µL of *P. aeruginosa* inoculum. The plates were incubated for 48 h at 37 °C, under aerobiosis and agitation (120 rpm). The culture medium was refreshed after 24 h.

### 2.4 CAP treatment of the biofilms formed on MGS

Biofilms were exposed to He-CAP treatment for 5 minutes. CAP was produced by the device described in Figure 1 under the same conditions employed to study the interaction between the plasma and the MGS, with a voltage amplitude of 12.3 kV p-p. For CAP treatment, each sample was placed exactly in the geometrical center of the well (from a 24-well plate), as seen in Figure 2.

**Figure 2.**
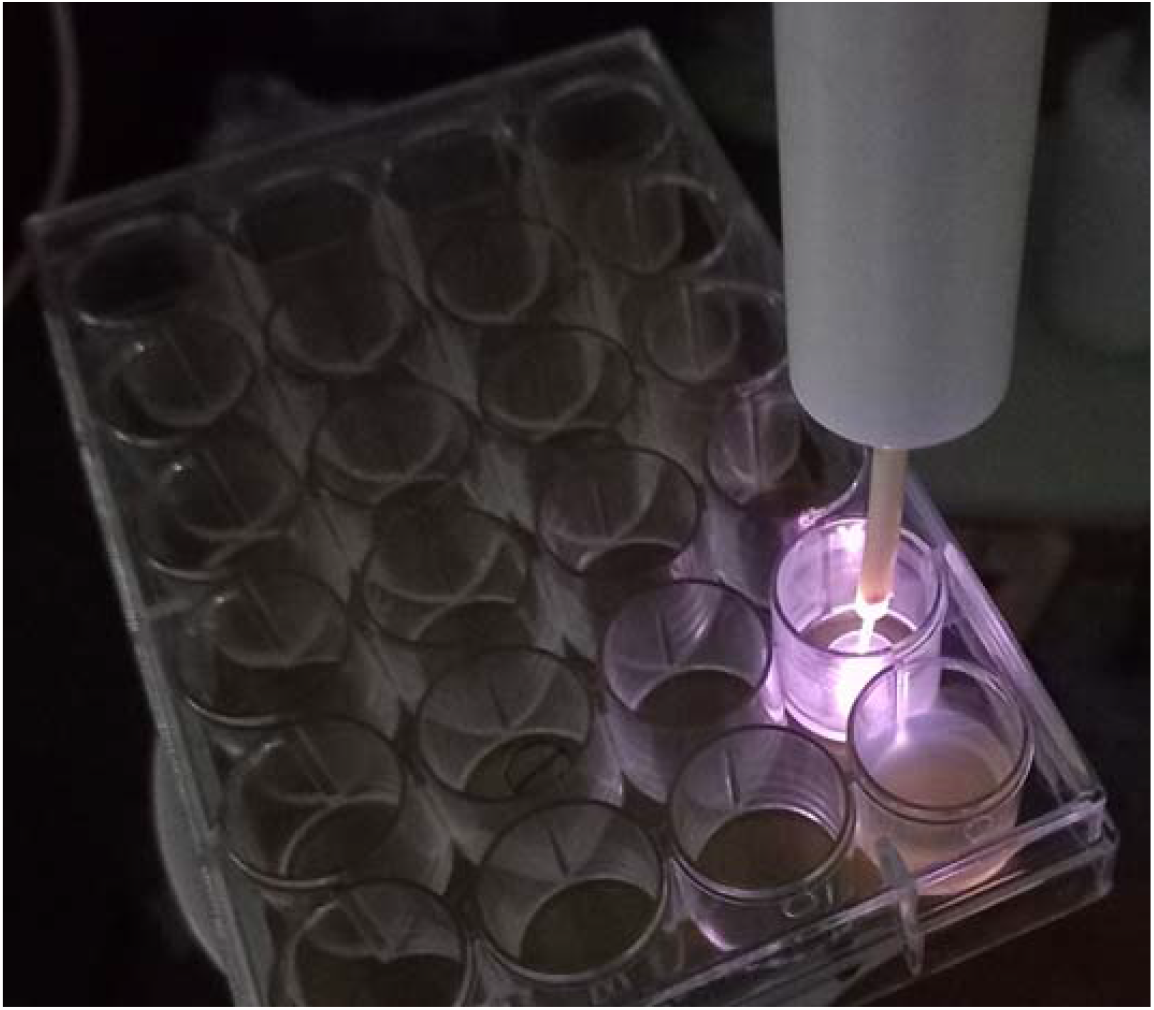
Application of the cold atmospheric plasma jet on the sample surface inside a well (24-well plate).

Biofilms were also flushed by He gas flow without plasma ignition for comparative purposes.

### 2.5 Determination of viable cell counts

After the CAP exposure, the biofilms were recovered from the MGS specimens by sonication (amplitude of 50%) in 3 cycles of 10 seconds pulse on, intercalated by 20 seconds pulse off. Finally, the suspensions were serially diluted and plated in specific agar to determine viable cell counts (CFU/mL), according to Table 2. The experiments were performed in triplicate on three independent occasions (n = 9). Data obtained were analyzed statistically using the Shapiro-Wilk normality test and compared using the Mann-Whitney test. The level of significance was set at 5%.

**Table 2.**
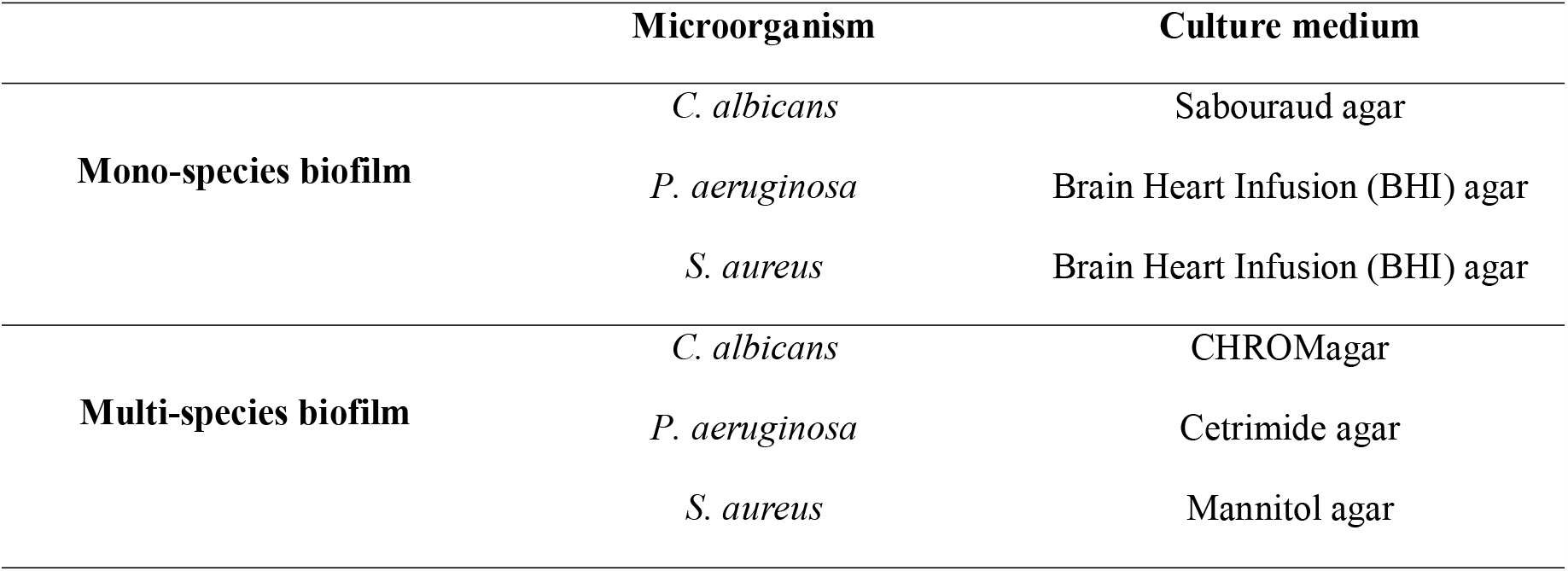
Description of each culture media used for viable cell recovery from mono and multi-species biofilms.

|  | Microorganism | Culture medium |
| --- | --- | --- |
| **Mono-species biofilm** | *C. albicans* | Sabouraud agar |
|  | *P. aeruginosa* | Brain Heart Infusion (BHI) agar |
|  | *S. aureus* | Brain Heart Infusion (BHI) agar |
| **Multi-species biofilm** | *C. albicans* | CHROMagar |
|  | *P. aeruginosa* | Cetrimide agar |
|  | *S. aureus* | Mannitol agar |

### 2.6 Cytotoxicity Analysis

The cell line BEAS-2B was selected to be used in this study to evaluate the toxicity of the CAP exposure protocol on the human bronchial epithelium. The cytotoxicity analysis was based on ISO 10993-5/2009 ^30^. Cells were incubated in Gibco ™ LHC-9 medium (Thermo Fisher) and kept incubated at 37 °C and 5% CO_2_ until they reached the desired confluence. To evaluate the cytotoxicity, 4 x 10^4^ cells per well were plated in 24-well plates. The experiments were performed in duplicate (n=12). For the treatment, 200 µL of the new medium was added to the wells to prevent them from drying out, and the cells were exposed to the products of CAP generated inside the T-tube.

Figure 3 shows the schematic for producing and applying the He CAP inside the tracheal T-tube. For this assay, the flexible tube from the CAP device was inserted horizontally inside the extratracheal portion. The 24-well plate was positioned under the T-tube vertical portion so that the distance between the CAP generated inside the T-tube and the cells was 2.0 cm. This configuration was used to study the effects of the CAP products on BEAS-2B cells. The treatment parameters were 31.7 kHz of frequency, 12.3 kV of voltage amplitude, and 2.0 SLM of helium gas flow, and the treatment group was exposed to CAP for 5 min. In the control group, the cells were not exposed to plasma.

**Figure 3.**
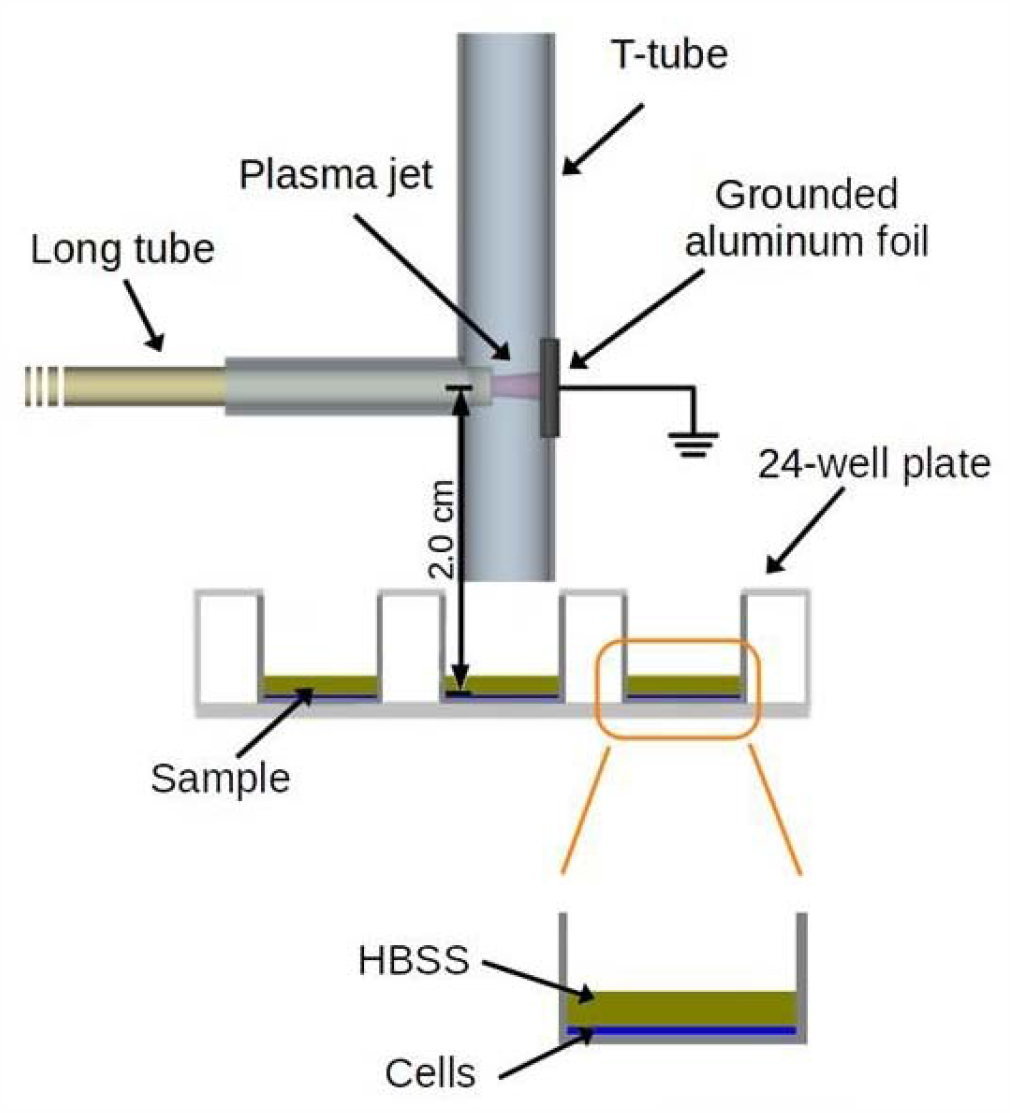
Schematic of the experimental setup to generate CAP inside the T-tube during the cytotoxicity test.

After CAP exposure of each well, 1 mL of the new medium was added to the wells. After 24 h, the medium was removed, the MTT reagent ((3-4,5-dimethylthiazol-2yl)-2,5-diphenyl-tetrazolium, Sigma, St. Louis, MI, USA) was added, and the plates were kept in agitation for 10 min. The optical density of the resulting solution was measured using a spectrophotometer at 570 nm. The control group normalized the obtained absorbance (=100%). Obtained data were evaluated in the GraphPad Prism software, version 8 (GraphPad Software, Inc, La Jolla, CA, USA). Cytotoxicity classification was based on the cell viability percentage, where values above 70% were considered non-cytotoxic ^31^.

## 3. RESULTS

### 3.1 MGS surface characterization after CAP exposure

The WCA behavior measured on the surface of the MGS samples after exposure to CAP treatment is presented in Figure 4. Images of the water droplets on the MGS samples are shown in Figure 5. Non-treated specimens have a WCA of 80.5°, i.e., exhibiting a less hydrophilic surface when compared to the control. The sample analyzed immediately after the CAP exposure showed a WCA < 5°, indicating that CAP changed the MGS surface properties to a highly hydrophilic surface. The MGS surface characteristics after CAP exposure that is, WCA < 5°, were maintained for approximately 15 min. After 30 min, the WCA increased to 15.2°, and this increasing trend was also observed for the sample analyzed after one hour, two hours, and four hours, showing WCA of 17.6°, 44.6°, and 53.2°, respectively.

**Figure 4.**
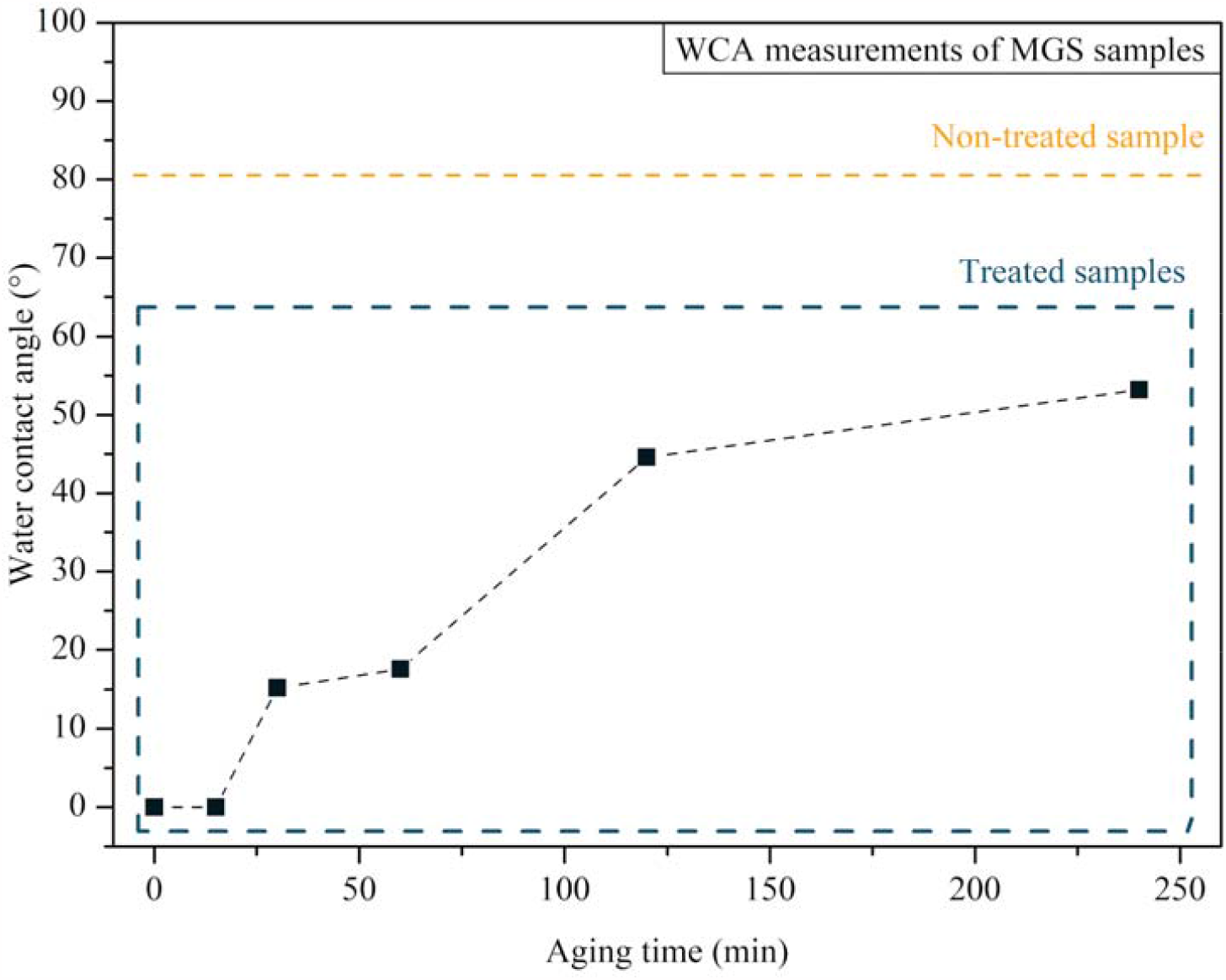
Aging time test of MGS samples after the treatment of CAP. The period of analysis varies between 0 min and 4 hours. The WCA of the non-treated sample is displayed in a box inside the graphic for comparison purposes.

**Figure 5.**
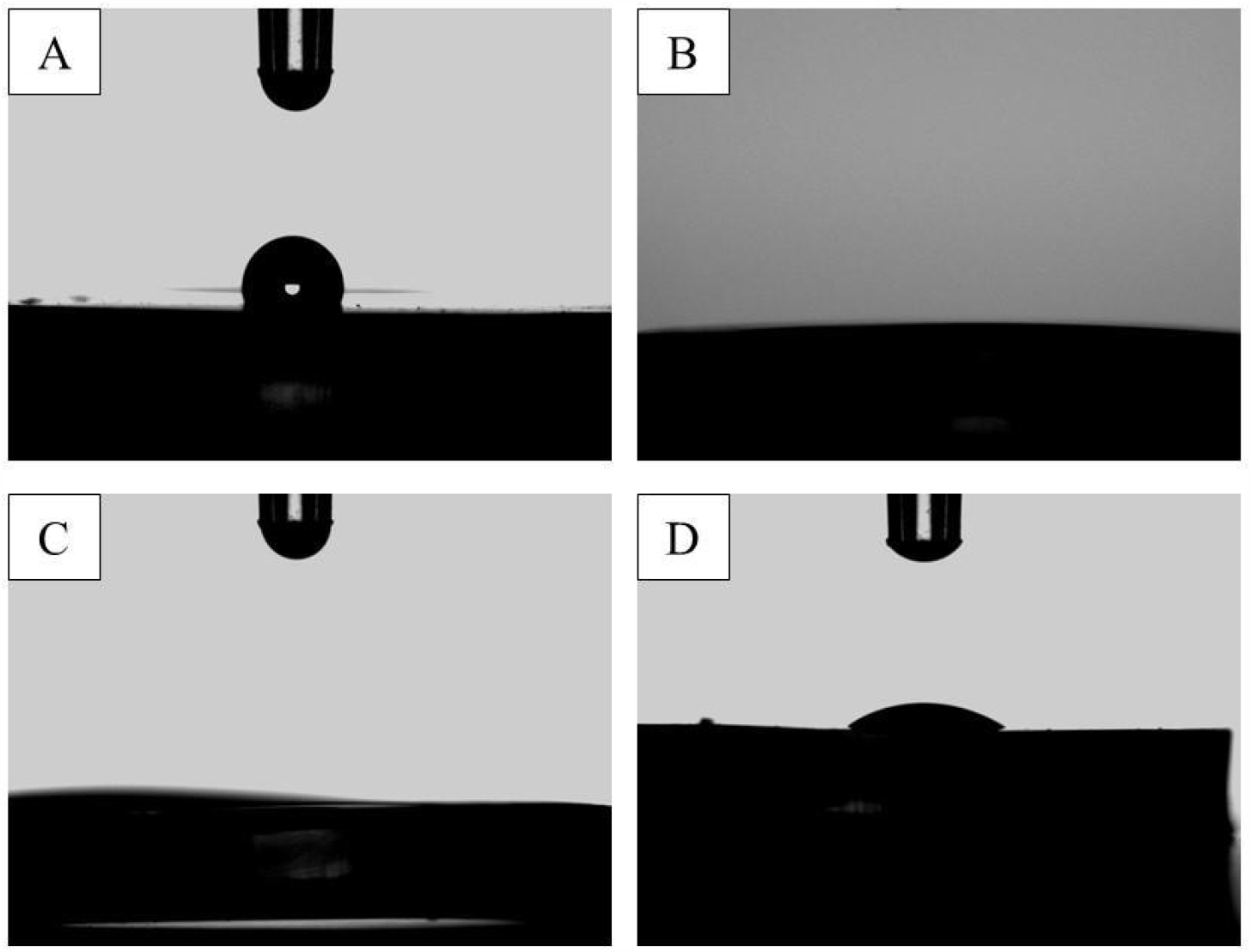
WCA measurement images showing the behavior of MGS surfaces in contact with water droplets at different times: (A) before CAP exposure; (B) Right after CAP exposure; (C) 15 min after CAP exposure; (D) 4 h after CAP exposure.

Figure 6 shows the spectra obtained from ATR-FTIR analysis of non-treated and treated-CAP samples. This analysis aimed to look for possible changes in the MGS compositional properties before and after the plasma irradiation. Table 3 contributes to the spectra analysis, presenting the correlation between the detected bands and the main functional groups on the MGS surface. The spectra of both samples demonstrate the C-H stretching band at 2962.3 cm^-1^, the asymmetric stretching of methyl groups at 1412 cm-1 characteristic for Si-CH_3_, and the symmetric stretching of the same group at 1258 cm^-1^. Finally, the typical bands at 800-1000 cm^-1^ are attributed to the Si-O-Si bond stretching ^32^.

**Table 3.** Attribution to the bands identified in the FT-IR analysis.

| Functional Group | Wavenumber ($\text{cm}^{-1}$) | Reference |
| --- | --- | --- |
| $-\text{CH}_3$ of $\text{SiCH}_3$ | 1258 and 1412 | 11 |
| Si-O-Si | 800-1000 |  |
| C-H | 2962.3 |  |

**Figure 6.**
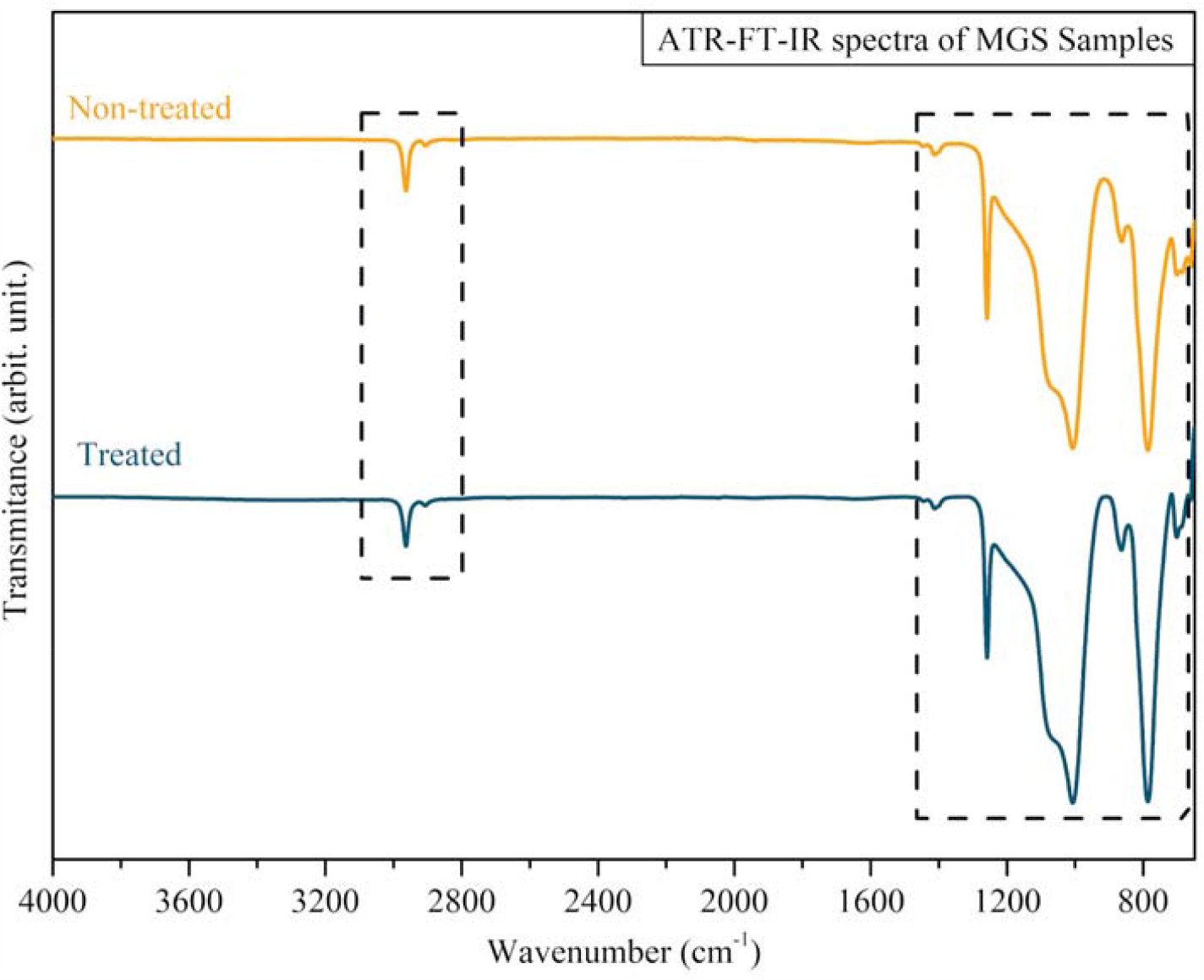
ATR-FTIR spectra of treated (blue) and non-treated (yellow) MGS samples.

### 3.2 Antimicrobial activity of CAP

The effect of CAP on mono-species biofilms of *C. albicans, P. aeruginosa*, and *S. aureus* formed on the MGS surface can be observed in Figure 7. *C. albicans* CAP treated group demonstrated an average count of viable cells of 1.362 x 10^6^ (CFU/mL), while the control group was 3.151 x 10^5^(CFU/mL). The treatment resulted in a statistically significant log reduction of 0.636 (T-test, p < 0.0001). The plasma-treated *P. aeruginosa* group presented an average count of viable cells of 1.42 x 10^6^ (CFU/mL), while the average value of the control group was 3.955 x 10^7^ (CFU/mL). In this way, there was a statistically significant log reduction of 1.445 (Mann-Whitney, p < 0.0001). *S. aureus* mono-specie biofilm count of viable cells of 5.194 x 10^6^ (CFU/mL) and 2.844 x 10^7^ (CFU/mL) were observed for treated and control groups, respectively, with a statistically significant log reduction of 0.738 (T-test, p < 0.05).

**Figure 7.**
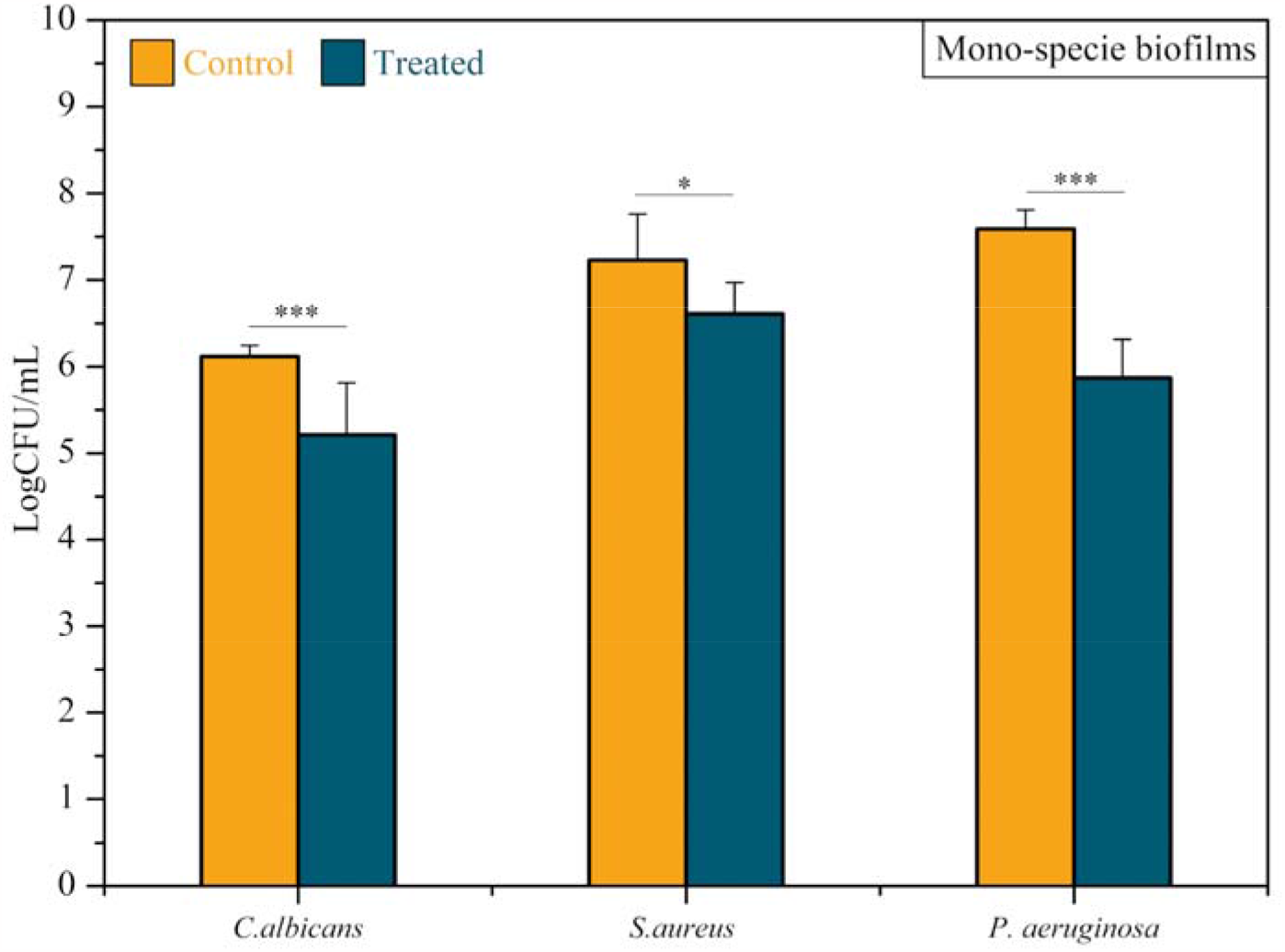
Antibiofilm efficacy of CAP on *Candida albicans, Staphylococcus aureus, and Pseudomonas aeruginosa* mono-specie biofilms. Results expressed in the logarithm of colony-forming units per milliliter (log CFU/mL). T-test *** p < 0.0001 and * p < 0.05.

Multispecies biofilms composed of *C. albicans, P. aeruginosa, and S. aureus* were also treated with CAP. Figure 8 shows the obtained results. The average count of *C. albicans* viable cells recovered from plasma-treated biofilm was 6.333 x 10^3^ (CFU/mL), while from multi-species biofilm control was 1.571 x 10^5^ (CFU/mL), indicating a statistically significant log reduction of 1.358 (Mann-Whitney test, p < 0.0001). Plasma-treated *P. aeruginosa* biofilms had 1.551 x 10^5^ (CFU/mL) viable cells, while 1.655 x 10^5^ (CFU/mL) viable cells were recovered from the control biofilm. Thus, *P. aeruginosa* showed a slight log reduction of 0.028 (p > 0.05). The average value of *S. aureus* viable cells recovered from the plasma-treated group and the control group was 1.0 x 10^3^ (CFU/mL) and 6.77 x 10^3^ (CFU/mL), respectively. There was a statistically significant log reduction of 0.83 (Mann-Whitney test, p < 0.0001).

**Figure 8.**
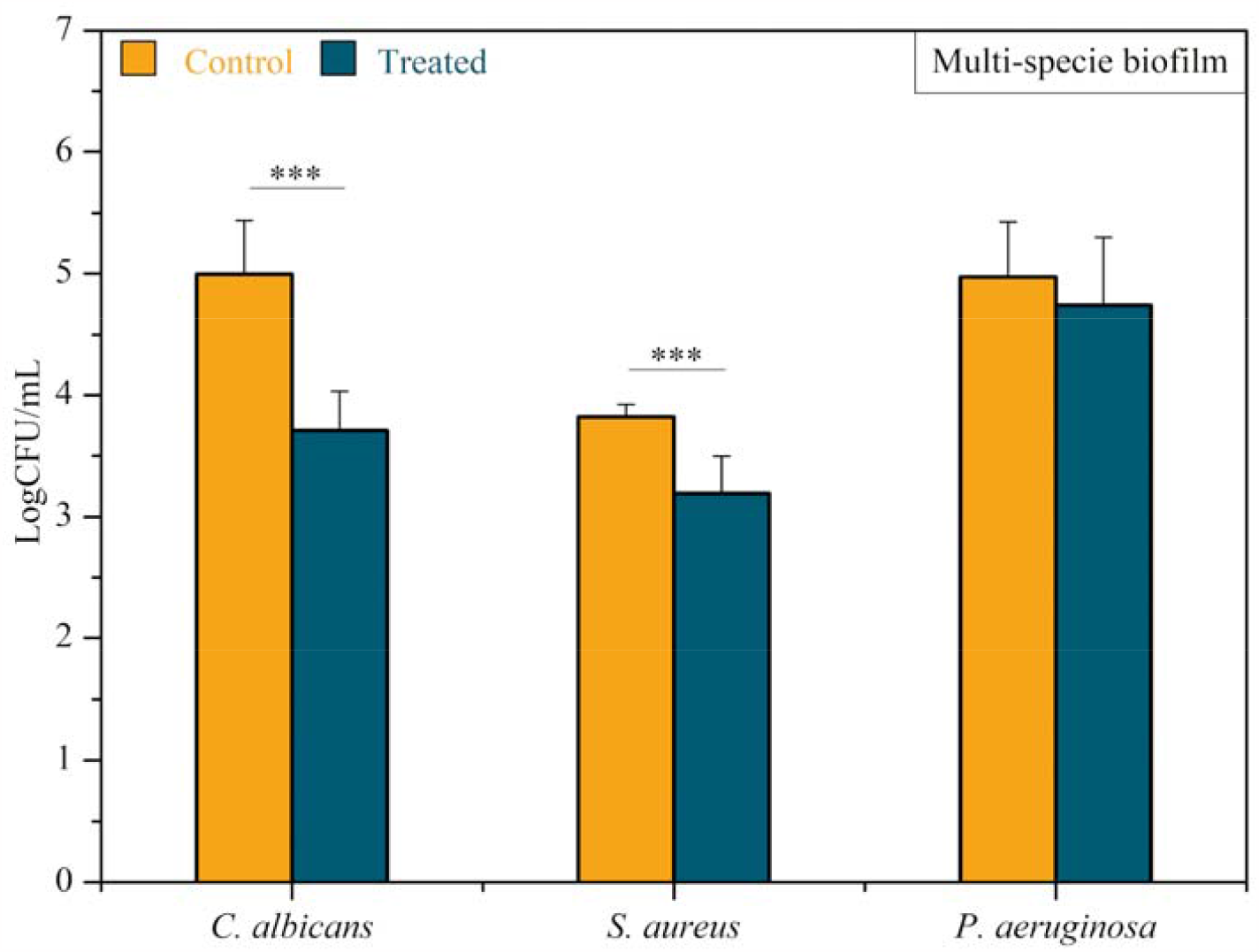
Antibiofilm efficacy of CAP on *C. albicans, P. aeruginosa, and S. aureus* multispecies biofilms. Results are expressed in the logarithm of colony-forming units per milliliter (CFU/mL). Mann-Whitney test, *** indicates p < 0.0001.

### 3.3 Cytotoxicity test

The results of the cytotoxicity test using the BEAS-2B cells are presented in Figure 9. The viability of cells exposed to CAP generated inside the T-tube was 95.76%. Thus, the protocol with antimicrobial activity can be considered non-cytotoxic (viability >70%).

**Figure 9.**
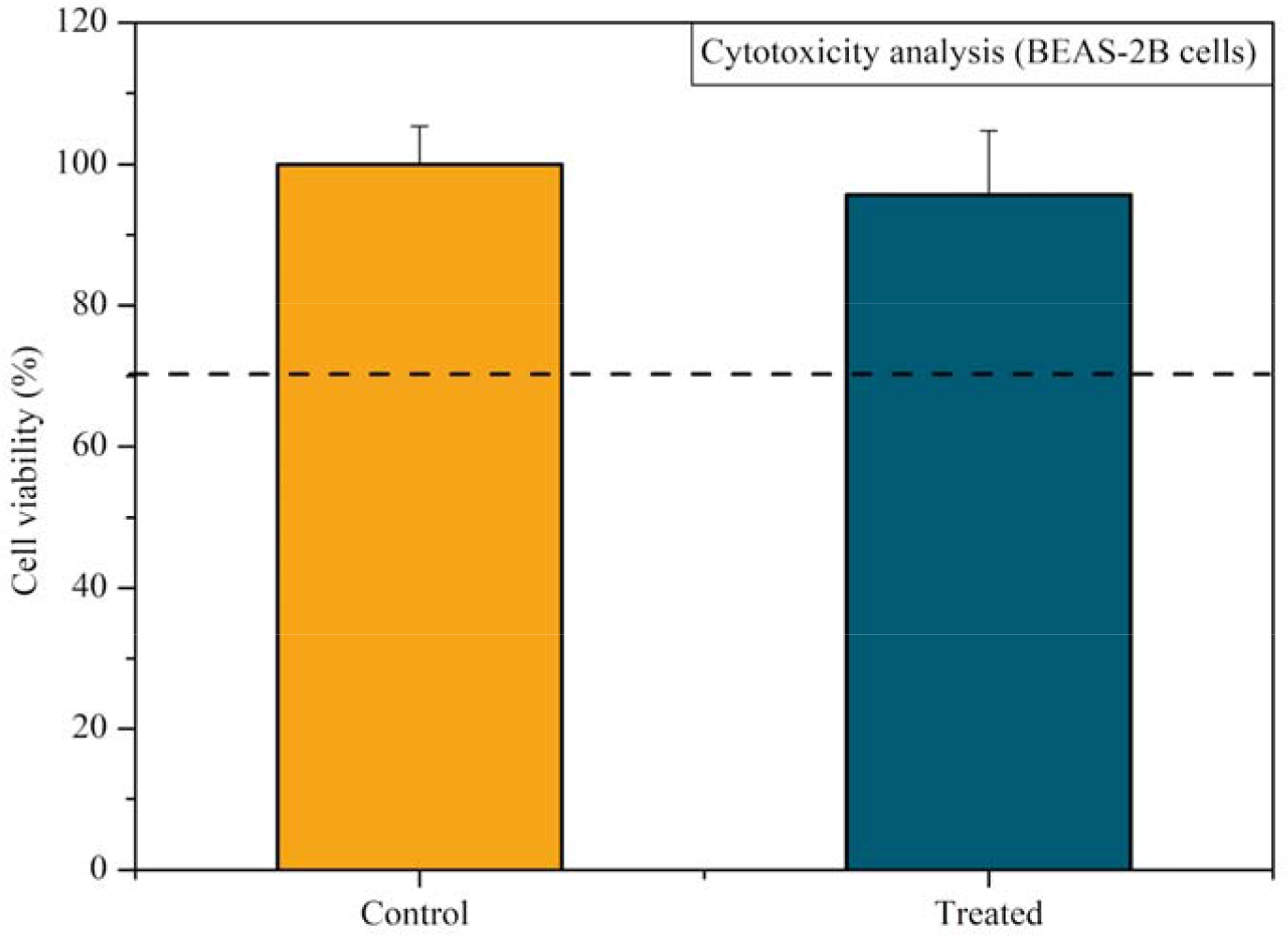
Cytotoxicity analysis of CAP expressed in cell viability (control 100%). The BEAS-2B cells were exposed to CAP generated inside the T-tube for 5 min. The cell viability was determined 24 h after CAP exposure. The dashed line represents the 70 % of cell viability.

## 4. DISCUSSION

The increase in tracheal stenosis cases related to previous episodes of COVID-19 emerged as an alarming reality after the pandemic. Beyouglu et al. (2022) reported that the intubation time of patients with COVID-19 is significantly longer when compared to patients with other conditions, which further increases the occurrence of tracheal stenosis and the subsequent use of silicone endotracheal prostheses ^33^. To avoid the prostheses changing every 6 to 12 months after implantation, alternative treatments that do not negatively interfere with the properties of MGS and present disinfection properties are needed. This study investigated CAP as an antimicrobial tool for removing biofilm commonly found in silicone endotracheal prostheses.

In the first stage, the MGS surface characterization was performed after CAP exposure. WCA increased steadily with time after the CAP treatment. Otherwise, in the period studied, it did not recover to the same WCA before CAP exposure (80.5°). Thus, the plasma treatment temporarily changes the MGS surfaces, turning them highly hydrophilic; in the middle, the WCA increases but does not reach the same contact angle as the non-treated sample. After the treatment, the MGS surface at 4 h is still more hydrophilic than the untreated MGS.

Regarding the FT-IR measurements, comparing both spectra, no significant differences were observed in the treated MGS sample compared to the untreated one. This is a positive result, as it shows that the plasma interacts with the MGS material, changing its surface energy without causing any change in its chemical composition or structure.

The hydrophobicity changes highlighted in the WCA analysis can be attributed to the interaction between the MGS surfaces and the RONS produced by CAP ^34^, leading to the formation of polar groups. Conversely, the ATR-FTIR analysis did not detect these plasma-induced polar groups. However, we should consider that when an internal reflection element is used in ATR-FTIR analysis, the IR beam detects chemical groups to the depth of 0.5-5 µm ^35^. So, suppose the plasma modification is limited only to the sample surface. In that case, the ATR-FTIR probably will not be able to detect any changes because it integrates the material response over a much broader depth. On the other hand, the behavior highlighted in the WCA aging test on the MGS surface occurs in the outermost surface layers (few nm) that were affected by the plasma. In this sense, further analyses more sensitive in detecting chemical changes on the MGS surfaces may clarify some of our findings.

To the best of our knowledge, the usage of CAP to treat biofilms on MGS surfaces at our set-up conditions has not been previously reported. However, biofilms treated with CAP on different substrates have already been investigated, and it will be a parameter to discuss our results.

In a previous study, S. aureus (ATCC 33591) and P. aeruginosa (ATCC 27853) biofilms formed on collagen membranes and were treated with CAP for 5 min with 15 mm of working distance. The treatment against these biofilms was as effective as the treatment performed in our study. The authors observed higher *S. aureus* log reduction than our experiments for the same microorganism ^36^. However, it is worth mentioning that we used a different *S. aureus* strain, which could justify some differences in the data. In this previous work, the reported log reduction for *P. aeruginosa* biofilm was lower than the decrease observed in our experiments for the same strain (ATCC 27853). These results indicate that reducing the working distance to 5 mm improves the antimicrobial effectiveness of *P. aeruginosa* biofilm.

The findings in the literature on CAP treatment against *C. albicans*, using similar parameters, are available to ATCC 18804. Inhibition zones of 2.0 and 2.5 cm were reached after 150 s of treatment ^23^. Another study with *C. albicans* strain SC 5314 reported a 2-log reduction when CAP was applied using the same parameters used in the present study and 15 mm of working distance^37^. It is well known that this strain is very virulent, reinforcing the antifungal potential of CAP treatment besides its antibacterial effect.

Our study also demonstrated the potential of CAP for treating multispecies biofilms, which more closely reflects the clinical situation of the endotracheal prostheses. *P. aeruginosa* was the only microorganism that did not show a statistically significant reduction after treatment of multispecies biofilms. This finding probably occurred due to *P. aeruginosa* dispersal in the polymeric extracellular matrices within the *C. albicans* and *S. aureus*, with no or little interaction between *P. aeruginosa* and CAP reactive species. Sequential applications of CAP can be an alternative to enhance the antibacterial activity against *P. aeruginosa* in multispecies biofilms. Once this study did not explore the sequential application of CAP, further studies are necessary better to understand its effects on P. aeruginosa in multispecies biofilms.

In addition to being antimicrobial, the proposed protocol was not cytotoxic to tracheal cells (BEAS-2B) in the conditions of this study. We emphasize that we focused on exposing the MGS used in T-tubes to CAP and not directly to the cells, simulating a clinical situation where the CAP jet will be directed to the contaminated prosthesis. We observed that a controlled application of CAP in the prosthesis is unlikely to cause harm to the epithelial cells adjacent to and outside the wall of the T-tube. Future studies are necessary to clarify the distribution of RONs inside the tube and to check the improvement of the antimicrobial activity after sequential exposures since promising findings have already been obtained after only one application.

## 5. CONCLUSION

In conclusion, CAP exposure of MGS samples enhances the surface hydrophilic properties from hydrophilic (WCA= 80.5°) to highly hydrophilic (WCA < 5°). The plasma jet-induced changes on the surface of the sample were not permanent. ATR-FTIR analysis did not show any significant differences between CAP-treated and non-treated samples, indicating essentially only a modification on the surface of the sample. CAP showed an effective inhibitory effect on all mono-species biofilms formed on MGS surfaces. It also demonstrated antimicrobial efficacy against *C. albicans* and *S. aureus* cells in the multi-species biofilm. The cytotoxicity test showed that the protocol is not cytotoxic for BEAS-2B cells when CAP is generated inside the T-tube. Future studies are required to understand better CAP potential and limitations in controlling the polymicrobial biofilms inside the endotracheal prosthesis and to make further improvements in the protocol. However, with the findings of this *in vitro* study, it is already possible to state that CAP is a suitable tool for the disinfection of MGS surfaces and, therefore, promising for the disinfection of silicone endotracheal prostheses.

## 6. ACKNOWLEDGMENTS

This work was supported by the São Paulo State Research Foundation (FAPESP) Grant numbers 2019/05856-7, 2021/02680-5; Coordination for the Improvement of Higher Education (CAPES); and the Brazilian National Council for Scientific and Technological Development Grant numbers 309762/2021-9, 308127/2018-8.

## REFERENCES

1. Piazza, C. et al. Long-term intubation and high rate of tracheostomy in COVID-19 patients might determine an unprecedented increase of airway stenoses: a call to action from the European Laryngological Society. European Archives of Oto-Rhino-Laryngology vol. 278 Preprint at 10.1007/s00405-020-06112-6 (2021).

2. Esteller-Moré, E. et al. Prognostic factors in laryngotracheal injury following intubation and/or tracheotomy in ICU patients. European Archives of Oto-Rhino-Laryngology and Head & Neck 262, 880–883 (2005).

3. Mattioli, F. et al. Post-intubation tracheal stenosis in COVID-19 patients. European Archives of Oto-Rhino-Laryngology 278, 847–848 (2021).

4. Alturk, A., Bara, A. & Darwish, B. Post-intubation tracheal stenosis after severe COVID-19 infection: A report of two cases. Annals of Medicine and Surgery 67, 102468 (2021).

5. Auchincloss, H. G. & Wright, C. D. Complications after tracheal resection and reconstruction: Prevention and treatment. Journal of Thoracic Disease vol. 8 S160–S167 Preprint at 10.3978/j.issn.2072-1439.2016.01.86 (2016).

6. Rorris, F. P. et al. Tracheal resection in patients post–COVID-19 is associated with high reintervention rate and early restenosis. JTCVS Tech (2023) doi:10.1016/j.xjtc.2023.01.006.

7. Ershadi, R., Rafieian, S., Sarbazzadeh, J. & Vahedi, M. Tracheal stenosis following mild-to-moderate COVID-19 infection without history of tracheal intubation: a case report. Gen Thorac Cardiovasc Surg 70, 303–307 (2022).

8. Montgomery, W. W. T-Tube Tracheal Stent. Arch Otolaryngol 82, 320–321 (1965).

9. Dumon, J.-F. A Dedicated Tracheobronchial Stent. Chest 97, 328–332 (1990).

10. Bibas, B. J. et al. Health-related quality of life evaluation in patients with non-surgical benign tracheal stenosis. J Thorac Dis 10, 4782–4788 (2018).

11. Ding, K., Wang, Y., Liu, S., Wang, S. & Mi, J. Preparation of medical hydrophilic and antibacterial silicone rubber via surface modification. RSC Adv. 11, 39950–39957 (2021).

12. Bibas, B. J. et al. Predictors for Postoperative Complications After Tracheal Resection. Ann Thorac Surg 98, 277–282 (2014).

13. Fusconi, M. et al. Is Montgomery Tracheal Safe-T-Tube Clinical Failure Induced by Biofilm? Otolaryngology–Head and Neck Surgery 149, 269–276 (2013).

14. Lee, J. M. et al. Biofilm accumulation on endotracheal tubes following prolonged intubation. J Laryngol Otol 126, 267–270 (2012).

15. Nouraei, S. A. R. et al. Bacterial Colonization of Airway Stents. Arch Otolaryngol Head Neck Surg 132, 1086 (2006).

16. Raveendra, N., Rathnakara, S. H., Haswani, N. & Subramaniam, V. Bacterial Biofilms on Tracheostomy Tubes. Indian Journal of Otolaryngology and Head & Neck Surgery (2021) doi:10.1007/s12070-021-02598-6.

17. Mazhar, K. et al. Bacterial Biofilms and Increased Bacterial Counts Are Associated with Airway Stenosis. Otolaryngology–Head and Neck Surgery 150, 834–840 (2014).

18. Chen, X., Ling, X., Liu, G. & Xiao, J. Antimicrobial Coating: Tracheal Tube Application. Int J Nanomedicine Volume 17, 1483–1494 (2022).

19. Duarte, S. & Panariello, B. H. D. Comprehensive biomedical applications of low temperature plasmas. Arch Biochem Biophys 693, 108560 (2020).

20. Bekeschus, S., Saadati, F. & Emmert, S. The potential of gas plasma technology for targeting breast cancer. Clin Transl Med 12, (2022).

21. Martusevich, A. K. et al. Cold Argon Athmospheric Plasma for Biomedicine: Biological Effects, Applications and Possibilities. Antioxidants 11, 1262 (2022).

22. Miebach, L., Poschkamp, B., van der Linde, J. & Bekeschus, S. Medical Gas Plasma—A Potent ROS-Generating Technology for Managing Intraoperative Bleeding Complications. Applied Sciences 12, 3800 (2022).

23. Kostov et al. Inactivation of Candida albicans by Cold Atmospheric Pressure Plasma Jet. IEEE Transactions on Plasma Science 43, 770–775 (2015).

24. Kostov et al. Study of Cold Atmospheric Plasma Jet at the End of Flexible Plastic Tube for Microbial Decontamination. Plasma Processes and Polymers 12, 1383–1391 (2015).

25. Borges, A. C. et al. Amplitude-modulated cold atmospheric pressure plasma jet for treatment of oral candidiasis: In vivo study. PLoS One 13, e0199832 (2018).

26. Bastin, O. et al. Optical and Electrical Characteristics of an Endoscopic DBD Plasma Jet. Plasma Med 10, 71–90 (2020).

27. Decauchy, H., Pavy, A., Camus, M., Fouassier, L. & Dufour, T. Cold plasma endoscopy applied to biliary ducts: feasibility risk assessment on human-like and porcine models for the treatment of cholangiocarcinoma. J Phys D Appl Phys 55, 455401 (2022).

28. Kondeti, V. S. S. K. et al. Long-lived and short-lived reactive species produced by a cold atmospheric pressure plasma jet for the inactivation of Pseudomonas aeruginosa and Staphylococcus aureus. Free Radic Biol Med 124, 275–287 (2018).

29. Mrochen, D. M. et al. Toxicity and virucidal activity of a neon-driven micro plasma jet on eukaryotic cells and a coronavirus. Free Radic Biol Med 191, 105–118 (2022).

30. Iso, I. 10993–5: 2009 Biological evaluation of medical devices—part 5: tests for in vitro cytotoxicity. International Organization for Standardization, Geneva (2009).

31. Jablonská, E., Kubásek, J., Vojtěch, D., Ruml, T. & Lipov, J. Test conditions can significantly affect the results of in vitro cytotoxicity testing of degradable metallic biomaterials. Sci Rep 11, (2021).

32. Ceresa, C. et al. Medical-grade silicone coated with rhamnolipid R89 is effective against Staphylococcus spp. Biofilms. Molecules 24, (2019).

33. Beyoglu, M. A., Sahin, M. F., Turkkan, S., Yazicioglu, A. & Yekeler, E. Complex Post-intubation Tracheal Stenosis in Covid-19 Patients. Indian Journal of Surgery 84, 805–813 (2022).

34. Liu, G. et al. Penetration effect of the kINPen plasma jet investigated with a 3D agar-entrapped bacteria model. Microchemical Journal 183, 107973 (2022).

35. >Tiernan, H., Byrne, B. & Kazarian, S. G. ATR-FTIR spectroscopy and spectroscopic imaging for the analysis of biopharmaceuticals. Spectrochimica Acta - Part A: Molecular and Biomolecular Spectroscopy vol. 241 Preprint at 10.1016/j.saa.2020.118636 (2020).

36. Oliveira, M. A. C. de et al. Inhibitory Effect of Cold Atmospheric Plasma on Chronic Wound-Related Multispecies Biofilms. Applied Sciences 11, 5441 (2021).

37. Borges, A. C. et al. Cold atmospheric pressure plasma jet modulates Candida albicans virulence traits. Clin Plasma Med 7–8, 9–15 (2017).

